# Dynamics of chromosomal target search by a membrane-integrated one-component receptor

**DOI:** 10.1101/2020.06.28.176644

**Authors:** Linda Martini, Sophie Brameyer, Elisabeth Hoyer, Kirsten Jung, Ulrich Gerland

**Affiliations:** Physics of Complex Biosystems, Technical University of Munich, James-Franck-Str. 1, 85748 Garching, Germany; Microbiology, Ludwig-Maximilians-University Munich, Großhaderner Str. 2-4, 82152 Martinsried, Germany

## Abstract

Membrane proteins account for about one third of the cellular proteome, but it is still unclear how dynamic they are and how they establish functional contacts with cytoplasmic interaction partners. Here, we consider a membrane-integrated one-component receptor that also acts as a transcriptional activator, and analyze how it kinetically locates its specific binding site on the genome. We focus on the case of CadC, the pH receptor of the acid stress response Cad system in *E. coli.* CadC is a prime example of a one-component signaling protein that directly binds to its cognate target site on the chromosome to regulate transcription. We combined fluorescence microscopy experiments, mathematical analysis, and kinetic Monte Carlo simulations to probe this target search process. Using fluorescently labeled CadC, we measured the time from activation of the receptor until successful binding to the DNA in single cells, exploiting that stable receptor-DNA complexes are visible as fluorescent spots. Our experimental data indicate that CadC is highly mobile in the membrane and finds its target by a 2D diffusion and capture mechanism. DNA mobility is constrained due to the overall chromosome organization, but a labeled DNA locus in the vicinity of the target site appears sufficiently mobile to randomly come close to the membrane. Relocation of the DNA target site to a distant position on the chromosome had almost no effect on the mean search time, which was between four and five minutes in either case. However, a mutant strain with two binding sites displayed a mean search time that was reduced by about a factor of two. This behavior is consistent with simulations of a coarse-grained lattice model for the coupled dynamics of DNA within a cell volume and proteins on its surface. The model also rationalizes the experimentally determined distribution of search times. Overall our findings reveal that DNA target search does not present a much bigger kinetic challenge for membrane-integrated proteins than for cytoplasmic proteins. More generally, diffusion and capture mechanisms may be sufficient for bacterial membrane proteins to establish functional contacts with cytoplasmic targets.

**Author summary:** Adaptation to changing environments is vital to bacteria and is enabled by sophisticated signal transduction systems. While signal transduction by two-component systems is well studied, the signal transduction of membrane-integrated one-component systems, where one protein performs both sensing and response regulation, are insufficiently understood. How can a membrane-integrated protein bind to specific sites on the genome to regulate transcription? Here, we study the kinetics of this process, which involves both protein diffusion within the membrane and conformational fluctuations of the genomic DNA. A well-suited model system for this question is CadC, the signaling protein of the *E. coli* Cad system involved in pH stress response. Fluorescently labeled CadC forms visible spots in single cells upon stable DNA-binding, marking the end of the protein-DNA search process. Moreover, the start of the search is triggered by a medium shift exposing cells to pH stress. We probe the underlying mechanism by varying the number and position of DNA target sites. We combine these experiments with mathematical analysis and kinetic Monte Carlo simulations of lattice models for the search process. Our results suggest that CadC diffusion in the membrane is pivotal for this search, while the DNA target site is just mobile enough to reach the membrane.

## Introduction

Bacteria are exposed to fluctuating environments with frequent changes in nutrient conditions and communication signals, but also life-threatening conditions such as environmental stresses and antibiotics [1]. To sense and adapt to changing environmental conditions, bacteria have evolved sophisticated signaling schemes, primarily based on one- and two-component systems [2,3].

Two-component signaling systems feature a sensor kinase and a separate response regulator, where the former is typically membrane-integrated while the latter diffuses through the cytoplasm to reach its regulatory target [3]. The majority of response regulators are transcription factors that bind to specific target sites on the genomic DNA to activate or repress transcription. Hence, a key step in the signal transduction pathway of these two-component systems is a DNA target search by a cytoplasmic protein. The target search dynamics of cytoplasmic transcription factors have been thoroughly studied in the past decades, triggered by early *in vitro* experiments indicating that the *Escherichia coli* Lac repressor finds its target site faster than the rate limit for three-dimensional (3D) diffusion [4]. Inspired by the idea that a reduction of dimensionality can lead to enhanced reaction rates [5], the experiments were explained by a two-step process, where transcription factors locate their target by alternating periods of 3D diffusion and 1D sliding along the DNA [6]. Compared to pure 3D diffusion, sliding increases the association rate by effectively enlarging the target size (“antenna” effect) [7–9]. These dynamics were later probed with single-molecule methods, both *in vitro* [10,11] and *in vivo* [12,13]. For the Lac repressor as a paradigmatic example of a low copy number cytoplasmic transcription factor, the *in vivo* timescale of the target search was found to be about one minute [12]. Further studies continued to add to the detailed understanding of this target search process, e.g. with respect to effects of DNA conformation [14], DNA dynamics [15], and macromolecular crowding [16].

One-component signaling systems, in contrast, combine sensory function and response regulation within one protein [2]. The subset of one-component systems that are both membrane-integrated sensors and DNA-binding response regulators face an extraordinary DNA target search problem: They must locate and bind to a specific site on the bacterial chromosome from the membrane. This is the case for one-component systems of the ToxR receptor family, which have a modular structure featuring a periplasmic sensory domain followed by a single transmembrane helix connected via a linker to a cytoplasmic DNA-binding domain [18]. In addition to ToxR, the main regulator for virulence in *Vibrio cholerae,* members of this receptor family include TcpP and TfoS in *V. cholerae* [19], PsaE in *Yersinia tuberculosis* [20] and the pH stress-sensing receptor CadC in *E. coli* [21]. *A priori,* it is not clear how these one-component systems establish functional protein-DNA contacts after stimulus perception. One scenario is that the DNA-binding domain is proteolytically cleaved, such that it can search for its specific binding site in the same manner as a regular cytoplasmic transcription factor. Another possible scenario is that simultaneous transcription, translation, and membrane-insertion (“transertion” [22, 23]) tethers the DNA locus of the one-component system to the membrane and, at the same time, places the protein in the vicinity of its binding site on the DNA (which is typically close to the gene encoding the one-component system). The third scenario is a diffusion and capture mechanism [24], whereby the one-component system diffuses within the membrane and captures conformational fluctuations that bring the DNA close to the membrane.

For the case of *E. coli* CadC, a well characterized model system [25], indirect evidence [26,27] paired with direct observation [28] argues against the proteolytic cleavage and transertion scenarios, and instead supports the diffusion and capture mechanism. For instance, the transertion mechanism should be sensitive to relocating the *cadC* gene to a locus far from its native position, which is close to the CadC target site at the *cadBA* promoter. However, relocation to the distant *lac* operon did not reduce the regulatory output of CadC. One of the observations arguing against proteolytic cleavage was that external signals can rapidly deactivate the CadC response after the original stimulus, whereas cleavage would irreversibly separate the DNA-binding domain from the signaling input [26]. In contrast, the diffusion and capture mechanism appeared consistent with experiments that imaged fluorophore-labeled CadC *in vivo* [28]. These experiments showed that localized CadC spots in fluorescent microscopic images form after cells were shifted to a medium that simultaneously provides acid stress and a lysine-rich environment, the two input signals required to stimulate the CadC response [26]. Formation of these spots was rapidly reversible upon removal of the input signals. Furthermore, the observed number of CadC spots was positively correlated with the number of DNA-binding sites, indicating that the spots correspond to CadC-DNA complexes with a much lower mobility than freely diffusing CadC in the membrane.

Taken together, the existing data suggest that the one-component system CadC establishes the protein-DNA contact required for transcription regulation not by a conventional target search akin to cytoplasmic transcription factors (Fig. 1A), but instead by 2D diffusion of the protein in the membrane and fluctuations of the DNA conformation that occasionally bring the DNA region of the target site close enough to the membrane to be captured by the protein (Fig. 1B). Intuitively, a successful diffusion and capture event seems highly unlikely. However, that such events occur was independently demonstrated in an experiment that artificially tethered the Lac repressor to the cell membrane [29]. This construct was indeed able to inducibly repress transcription from a chromosomal reporter. Hence, the striking questions are how this type of target search is kinetically feasible and on which timescale. Here, we address these questions using *E. coli* CadC as a model system. We measure the search time of CadC to its target DNA-binding site in single cells by probing the formation of fluorescent CadC spots at different times. This yields experimental search time distributions that we compare against so-called ‘first passage time' distributions obtained from stochastic models [30]. We also measure the mobility of a chromosomal locus in our experimental setup using a fluorescent repressor/operator system to inform kinetic Monte Carlo simulations of the target search dynamics. These simulations are able to reproduce the experimental behavior and to elucidate properties of the search process that we cannot obtain experimentally.

**Fig 1.**
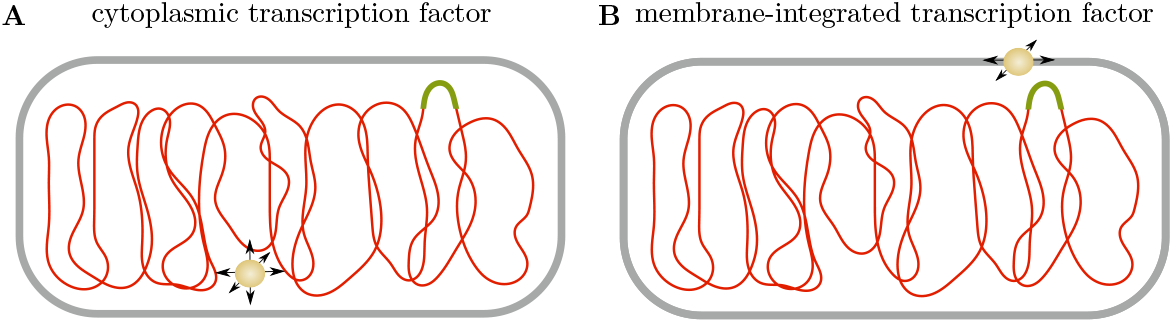
DNA target search of cytoplasmic vs. membrane proteins. **(A)** A cytoplasmic transcription factor (yellow) locates its specific binding site (green) on the DNA (red) through a combination of 3D diffusion in the cytoplasm and 1D sliding along the polymer. While the DNA itself is not static during this target search, DNA motion does not significantly contribute to its completion. **(B)** In contrast, a membrane-integrated transcription factor can only perform 2D diffusion in the membrane, such that DNA motion becomes essential. At a minimum, the specific DNA-binding site has to move close to the membrane to enable target recognition.

## Results

### Choice of CadC as experimental model system

CadC is a particularly well studied membrane-integrated one-component receptor. It is part of a pH stress response system, which is also dependent on signaling input from the lysine-specific permease LysP [26]. The function of the Cad system is to alleviate acidic stress by activating synthesis of the lysine/cadaverine antiporter CadB and the lysine decarboxylase CadA, which converts lysine to cadaverine to be secreted by CadB. CadA and CadB are encoded by the *cadBA* operon, which is transcriptionally upregulated by specific binding of CadC upstream of the promoter [31], see Fig 2A (inset). CadC is known to respond to and integrate three different signals: It is activated by an acidic pH in the presence of external lysine and inhibited by cadaverine. The pH-sensory function as well as the feedback inhibition by cadaverine was assigned to distinct amino acids within the periplasmic sensory domain of CadC [27, 32], whereas the availability of external lysine is transduced to CadC via the co-sensor and inhibitor LysP, a lysine-specific transporter [33,34]. High-affinity DNA-binding of CadC requires CadC homodimerization, which is inhibited by LysP via intramembrane and periplasmic contacts under non-inducing conditions [33, 34]. A drop in external pH induces dimerization of the periplasmic sensory domain of CadC followed by structural rearrangement of its cytoplasmic linker, permitting the DNA-binding domain of CadC to homodimerize [21,32,35]. The CadC protein number is extremely low (on average 1-3 molecules per cell [28]), mainly due to a low translation rate caused by polyproline stalling, which is only partially relieved by elongation factor P [36].

**Fig 2.**
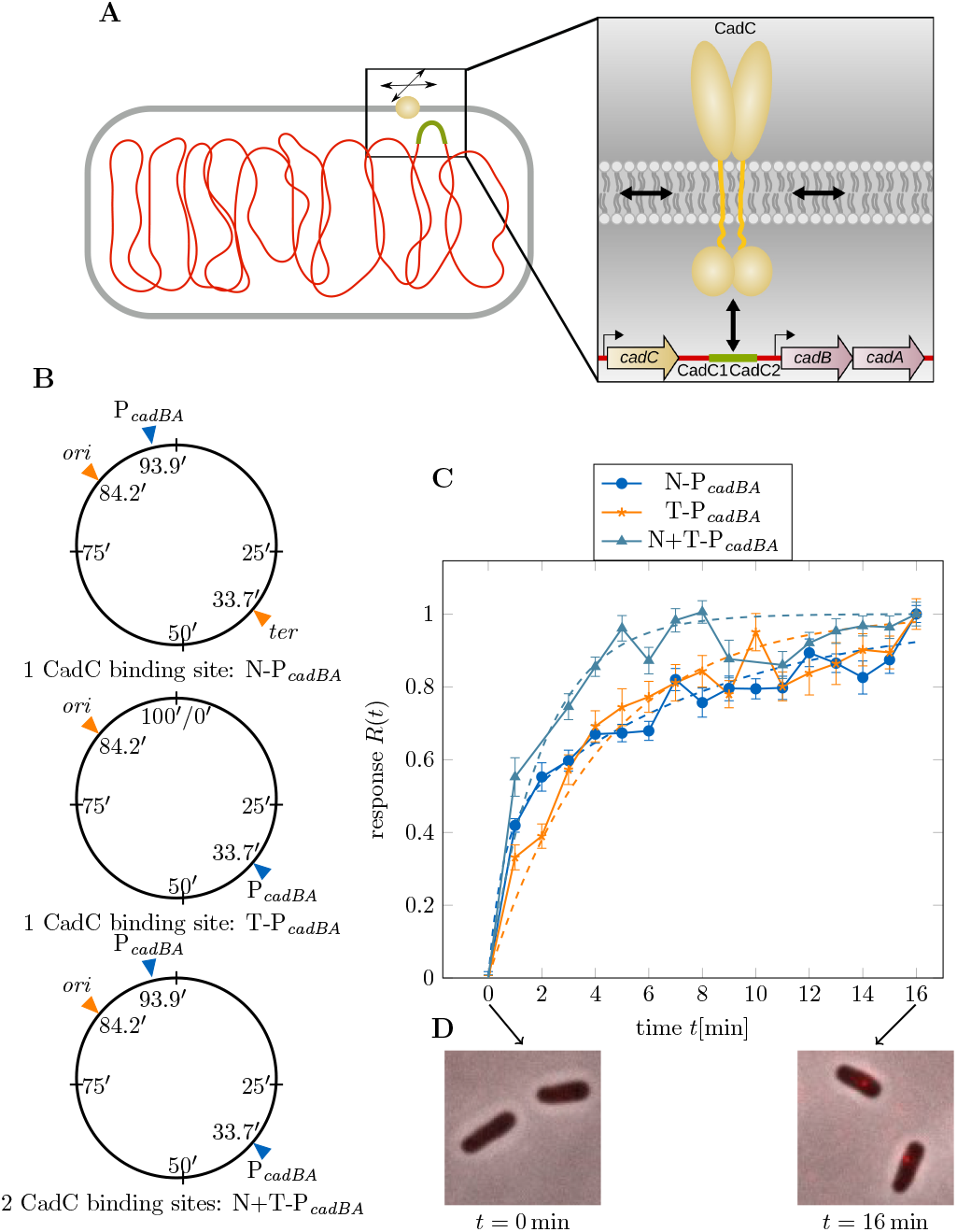
Dynamics of the target search by CadC. (**A**) The target search of membrane-integrated transcription factors is investigated by experimentally measuring the response time of CadC. The molecular model on the right shows CadC in yellow, diffusing in the membrane and forming two dimers to bind to the two CadC binding sites within the *cadBA* promoter (CadC1 and CadC2), displayed in green. The *cadC* gene is located upstream of the *cadBA* operon. (**B**) Three *E. coli* strains with different positions of the *cadBA* promoter: N-P*cadBA* with the native DNA-binding site close to *ori,* T-P_*cadBA*_ with the binding site at the terminus and N+T-P _*cadBA*_ with both binding sites. (**C**) Experimental results from CadC spot detection. Fluorescent microscopic images were taken every minute after receptor activation and analyzed for CadC spots for the strains defined in panel B. The plot shows the response *R*(*t*), defined as the normalized fraction of cells with fluorescent spots as a function of time t after exposure to acid stress. Error bars correspond to the propagated standard deviation of v(t) from averaging over multiple data sets. The dashed lines in the plot show a fit of the response function to the CDF of a sequential reversible two-step model with mixed initial condition for N-P _*cadBA*_, shown in blue dots and with fixed initial condition for T-P_*cadBA*_ (orange stars) and N+T-P_*cadBA*_ (cyan triangles). (**D**) The fluorescence microscopy images demonstrate how fluorescent spots appear after receptor activation.

In the present study, CadC serves as an experimental model system to investigate the DNA target search of a membrane-integrated transcription factor (Fig 2A). To probe the kinetics of this search process in individual cells, it is crucial to have a well-defined initial state and clear ‘start’ and ‘stop’ events. When cells are initially grown in a medium with neutral pH, CadC is inactive and homogeneously distributed in the membrane [28]. A sudden medium shift to low pH and rich lysine conditions then serves as a suitable ‘start’ trigger, which activates CadC via the signal transduction mechanism described above and starts the target search process. However, detecting the successful termination of the target search is challenging. Here, we exploit the previous finding that the formation of stable CadC-DNA complexes is visible as distinct spots in fluorescence microscopy images [28]. The study excluded that spot formation occurs solely due to low pH, using a pH-independent CadC variant that showed spots at both neutral and low pH. The connection between spot formation and CadC DNA-binding was derived from the observation that no spots were formed when CadC was rendered unable to bind DNA, and the number of spots per cell correlated with the number of DNA-binding sites for CadC. Without DNA-binding site only 20 % of cells formed spots, possibly due to non-specific DNA-binding. We take this into account in our quantitative analysis below.

### Experimental measurement of CadC target search times

We used three *E. coli* strains with different binding site configurations along the chromosome: N-P_*cadBA*_ (wild type) with the native DNA-binding site close to *ori* (at 93.9' [37]), T-P_*cadBA*_ with the binding site relocated to the terminus, and N+T-P_*cadBA*_ with both binding sites (Fig 2B). To visualize the temporal and spatial localization of CadC *in vivo*, we transformed each of the strains with plasmid-encoded mCherry-tagged CadC, thereby increasing the average number of CadC molecules per cell to 3-5 [28]. After the medium shift at *t* = 0 min, we took fluorescence and phase contrast microscopy images of cells sampled from the same culture every minute. We used image analysis tools to detect fluorescent spots within the cells, evaluating between 859 and 2506 cells per time step, see ‘Materials and methods’. Based on these data, we determined the fraction of cells with at least one fluorescent spot, *ν*(*t*) = *N*_cells with spot_(*t*)/*N*_cells_(*t*) at each time *t* after CadC activation. To take into account the initial fraction of spots attributed to non-specific DNA-binding (see above), we defined the response function

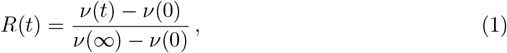

which rises from zero to one. Here, the asymptotic value *ν*(∞) accounts for the fact that fluorescent spots are never detected in all cells (see the raw data in S1 Fig). This is likely due to the heterogeneous distribution of CadC [28]: Given the low average copy number, some cells are expected to have less than the two molecules required for dimerization. Additionally, some spots may have been missed by the spot detection algorithm, in particular for cells that were not perfectly in focus. The time-dependent response *R*(*t*) for our three strains is shown in Fig 2C, with examples of fluorescence microscopy images of cells at *t* = 0min and the last time point (Fig 2D). Before analyzing the experimental response functions, we discuss the description of the target search as a stochastic process and derive theoretical response functions for comparison with the experimental data.

### The target search as a stochastic process

On a coarse-grained level, the search of CadC for a specific binding site on the DNA can be described by a stochastic process with a small number of discrete states. Since both CadC and the DNA must move in order to establish a specific protein-DNA contact, it is reasonable to assume a reversible sequential process with an intermediate state,

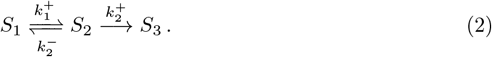

Here, state *S*_1_ corresponds to configurations where the DNA target site is not in direct vicinity of the membrane, and CadC is delocalized on the membrane, unbound to the DNA. State *S*_3_ corresponds to the final state where CadC is bound to a specific target site on the DNA. The intermediate state *S*_2_ could then correspond to configurations where the DNA segment containing the target site is close to the membrane, but CadC is not bound to this segment. The transition rates between these states are denoted as 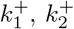, and 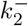. We are interested in the so-called ‘first-passage time’ *τ* to reach the final state S3, which corresponds to the target search time within this coarse-grained description. The probability distribution *p*(*τ*) for this time is calculated by making the final state absorbing, using standard techniques [30] (see S1 Appendix). Assuming that the system is initially in state *S*_1_, the first passage time distribution is

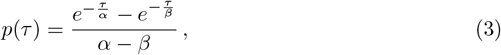

where the two timescales *α* and *β* of the exponential functions are related to the transition rates via

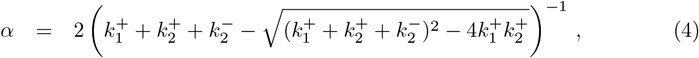

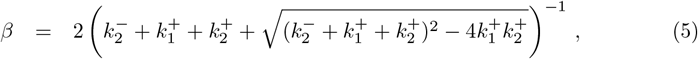

implying that *α* > *β*. At large times, the distribution *p*(*τ*) decays exponentially (decay time *α*), whereas the timescale ? corresponds to a delay at short times. Hence, increasing *α* leads to a slower decay and increasing *β* to a longer delay. The average value of this distribution, referred to as the mean first passage time, is

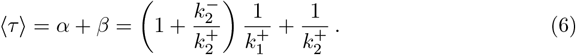

This corresponds to the average time for the first step 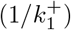 multiplied by the average number of trials needed to reach the second step, plus the average time for the second reaction step 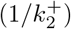. In cases where one of the reaction steps is rate limiting, the mean first passage time is simply the inverse of the limiting rate, and the process simplifies to a two-state process with an exponentially distributed first passage time.

To relate the first passage time distribution to our experimental response function, Eq 1, we consider the cumulative distribution function CDF(*τ*) defined by

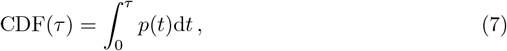

which is the probability that the first passage time is less than or equal to *τ*. Experimentally, CDF(*τ*) corresponds to the fraction of cells in which the target search was successful by time *τ*. The response function in Eq 1 is our best proxy for this fraction of cells, and hence we identify

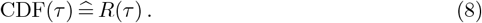

The cumulative distribution function for *p*(*τ*) of Eq 3 is

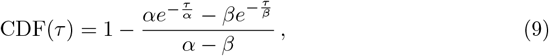

which we can use as a fit function to describe the experimental data. However, using Eq 9 amounts to the assumption that all cells are initially in state *S*_1_. If we allow for the possibility that some cells are in state *S*_2_ when CadC is activated, we have a mixed initial condition, where the process starts either from state *S*_1_ or from state *S*_2_. Denoting the fraction of cells that are initially in state *S*_2_ by *x*, such that a fraction

**Table 1.** Moments of the measured search processes.

| Strain | $\langle \tau \rangle [\text{min}]$ | $\sigma^2 [\text{min}^2]$ |
| --- | --- | --- |
| N-P<sub>cadBA</sub> | $4.84 \pm 0.19$ | $49.6 \pm 8.8$ |
| T-P<sub>cadBA</sub> | $4.20 \pm 0.15$ | $17.6 \pm 2.0$ |
| N+T-P<sub>cadBA</sub> | $2.02 \pm 0.12$ | $4.09 \pm 0.66$ |

Results from fitting the experimentally computed CDFs to the sequential reversible model with mixed initial condition (N-P<sub>cadBA</sub>) and fixed initial condition (T-P<sub>cadBA</sub> and N+T-P<sub>cadBA</sub>). The fit parameters $\alpha$ , $\beta$ and $c$ were used to compute the mean first passage time $\langle \tau \rangle$ and the variance $\sigma^2$ with uncertainties obtained from error propagation using the full covariance matrix.

1 – *x* starts in state *S*_1_, we obtain a first passage time distribution and associated cumulative distribution function of the form

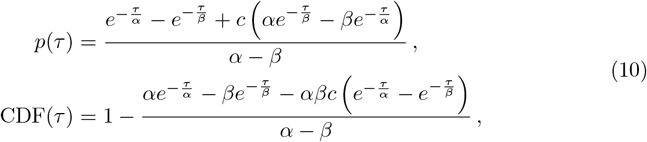

where the new parameter c corresponds to 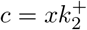. The associated mean first passage time is

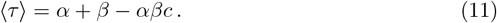

### The mean search time is less than 5 min and not affected by relocation of the target site

We first analyzed the experimental response *R*(*t*) in the wild type strain. The wild type data (N-P_*cadBA*_) in Fig 2C (blue dots) show an initially fast increase, followed by a more gradual saturation. This behavior is not well described by Eq 9, but captured by Eq 10 with the mixed initial condition, as can be seen by the fit to the data represented by the blue dashed line in Fig 2C. From the fit parameters *α, β*, and *c*, we computed the mean search time according to Eq 11, finding 〈*τ*〉 ≈ 4.80 ± 0.19 min. This result is consistent with the transcriptional response of the target genes *cadBA*, which was previously probed by Northern blot analysis, finding that the cell-averaged *cadBA* mRNA level starts to increase about 5 min after receptor activation [26].

In order to assess whether the position of the DNA-binding site along the chromosome affects the target search, we analyzed the behavior of strain T-P_*cadBA*_, which has the CadC DNA-binding site at the terminus instead of the native position. The corresponding response function shown in Fig 2C (orange stars) features a less pronounced initial increase than observed for N-P_*cadBA*_, and the dashed orange line shows an adequate fit using the sequential model with fixed initial condition in Eq 9. The calculated mean first passage time of 〈*τ*〉 ≈ 4.20 ± 0.15 min is comparable to that for the wild type strain. The slower initial increase is compensated by a faster increase at later times to yield a slightly smaller mean search time. To characterize the shape of the mean first passage time distributions, we calculated the variance *σ*^2^ of *p*(*τ*), see Table 1, which is smaller for strain T-P_*cadBA*_ than for the wild type.

### Search time is decreased with two chromosomal CadC binding sites

After observing essentially the same mean search time for two very distant locations of CadC target sites on the chromosome, we wondered how a strain harboring both target sites would behave. We therefore repeated the measurements for *E. coli* strain N+T-P_*cadBA*_, which has the native DNA-binding site and additionally the binding site at the terminus. As shown in Fig 2C (cyan triangles), the response function of this strain saturates much earlier than for the other two strains. Fitting the response data to the sequential model with fixed initial condition (Eq 9), we obtained a mean first passage time of 〈*τ*#x232A; ≈ 2.02 ± 0.12 min, which is only around half of the time than for a single chromosomal binding site.

### Colocalization of CadC spots with the DNA-binding site

We also wondered whether the fluorescence spots indicating the position of stable CadC-DNA complexes in single cells would show a similar spatial distribution as the *cadBA* locus. We therefore analyzed the localization of CadC spots along the long axis of the cell in *E. coli* wild type. As an estimate for the position of the *cadBA* locus along the cell, we tracked the position of *ori* at low pH. Towards this end we inserted a *parS* gene close to *ori* and let ParB-yGFP bind to it, making the *ori* region visible as a fluorescent spot in microscopy images. As the position of chromosomal loci depends on the progression of the cell cycle [38], we grouped cells according to their length into three classes. Fig 3 shows the spatial distribution of the relative spot positions along the half long axis for these three length classes, comparing ParB spots (blue) to CadC spots (orange). The large overlap of the distributions implies a similar cell age dependent localization of *ori* and CadC spots along the long cell axis, suggesting that CadC spots indeed form close to the DNA-binding site.

**Fig 3.**
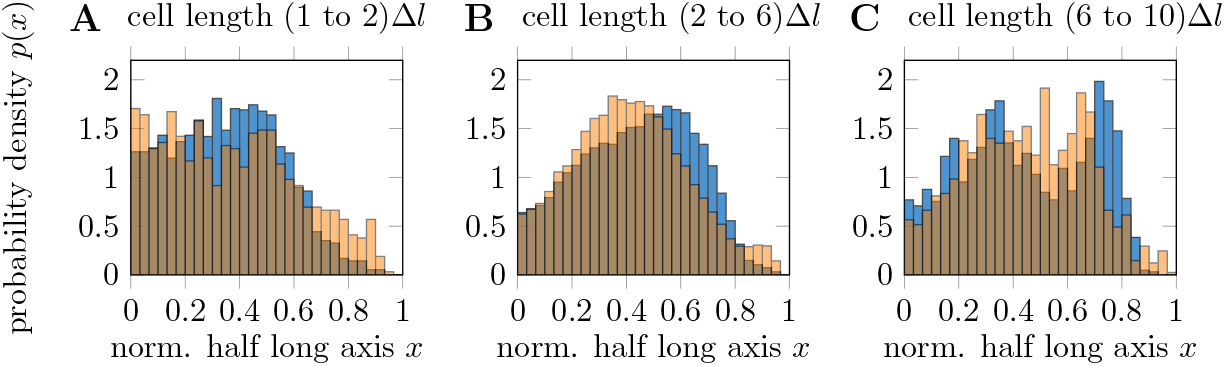
Localization of *ori* and CadC spots. Localization probability of CadC spots in N-P _*cadBA*_ cells (orange) and ParB localization marking *ori* (blue) along the half long axis of cells. The half long axis is normalized such that mid-cell is at *x* = 0 and the poles are at *x* = 1. Overlaps of the two distributions are shown in darker orange. Cell age is taken into account by splitting all occurring cell lengths into ten equally spaced steps Δ*l* and pooling the cells according to their size. From the ten different age classes we observed similar localization probabilities for *l* = (1 to 2)Δ*l* (**A**), *l* = (2 to 6)Δ*l* (**B**) and *l* = (6 to 10)Δ*l* (**C**), which are therefore grouped together in this plot.

### Biophysical model of the target search process

To gain more insight into the dynamics of the target search, we turned to a coarse-grained biophysical model for the coupled dynamics of CadC and the DNA. We simulated the search of a CadC dimer for its target DNA-binding site using a lattice model and a kinetic Monte Carlo approach. As depicted in Fig 4A, the DNA is represented by a path on the 3D lattice (the ‘cytoplasm’) and the CadC dimer moves on the surface of the lattice (the ‘membrane’). Starting from a random initial configuration, the CadC dimer diffuses in the membrane with a rate *k*_2D_, binds non-specifically to DNA segments that are at the membrane and close to CadC with rate *k*_on_ and unbinds (*k*_off_) or slides (*k*_1D_) along DNA segments at the membrane when bound. The simulation continues until the CadC dimer reaches the specific DNA-binding site to end the search process. As demonstrated in Fig 4B the DNA dynamics was implemented by random displacements of single beads, at a basal rate *k*_DNA_, with a kinetic Monte Carlo algorithm to accept moves according to a spring potential between neighboring beads, see ‘Materials and methods’. Similarly, CadC dimers move by random displacements constrained either to the membrane or to the polymer. For each set of parameters we run ≥ 10^3^ simulations to compute mean values and search time distributions.

**Fig 4.**
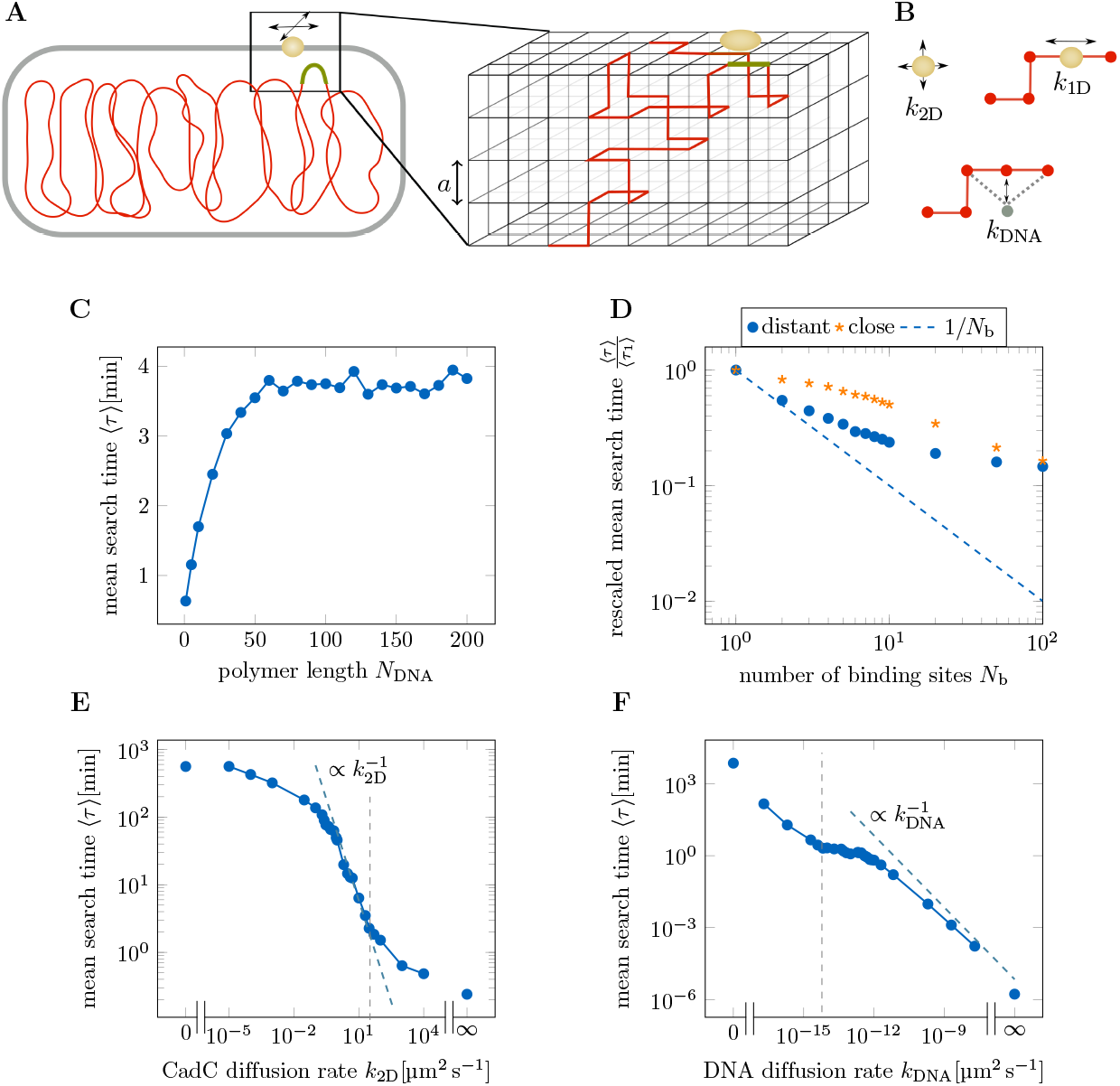
Characterization of the target search from computer simulations. (**A**) The target search of membrane-integrated transcription factors is studied by kinetic Monte Carlo simulations of a lattice model, shown on the right. The cubic lattice represents a cell with the DNA (red) inside and a CadC dimer (yellow) in the membrane. (**B**) The DNA moves by random displacements of single beads with rate *k*_DNA_. CadC dimers diffuse in the membrane with rate *k*_2D_, bind and unbind non-specifically to DNA segments at the membrane with rates *k*_on_, *k*_off_ and slide along DNA segments with rate *k*_1D_ when bound. (**C**) Mean target search time 〈*τ*〉 as a function of DNA length *N*_DNA_ using realistic parameters. (**D**) Dependence of the mean search time 〈*τ*〉, normalized by the search time with a single binding site 〈*τ*〉_1_, on the number of DNA-binding sites *N*_b_. Realistic parameters are used for binding sites placed either uniformly (blue dots) or next to each other (orange stars) along the circular chromosome. (**E**) Mean target search time for different CadC diffusion rates. The data point on the very left corresponds to *k*_2D_ = 0 μm^2^ s^-1^, the data point on the very right corresponds to *k*_2D_ → ∞ μm^2^ s^-1^, simulated by making the whole cell surface the target. The dashed gray line marks *k*_2D_ = 0.20 μm^2^ s^-1^, used in the realistic parameter set. (**F**) Mean target search time for different DNA diffusion rates. The dashed gray line marks *k*_DNA_ = 0.0060 μm^2^ s^-1^, used in the realistic parameter set.

To compare our simulations to the experiments we defined a parameter set based on known experimental estimates for most parameters, as discussed in ‘Materials and methods’ and summarized in Table 2. Instead of simulating the full length of an *E. coli* chromosome, we made use of an observed relation between search time and polymer length to reduce computation time. As shown in Fig 4C, the simulated mean search time first increases with the number of DNA segments, but then becomes independent of polymer length and reaches a plateau. We found both the plateau value and the critical polymer length at which it is reached to depend on the size of the simulated cell. Polymer segments far apart along the polymer become uncorrelated in their motion when the polymer in between touches the boundary. Therefore the confinement leads polymer subchains to become independent when the DNA is long enough compared to the simulated cell to frequently encounter the boundary. Hence it is not necessary to simulate the full length of an *E. coli* chromosome, as long as the simulated polymer length lies within the plateau region. For the realistic parameter set used in Fig 4C, simulating a polymer length of *N_DNA_* = 100 beads is sufficient, which is the minimum value we used in our simulations. To start with the simplest model, non-specific binding and sliding were not included in the realistic parameter set.

### Mobility measurements of the DNA

To complete our parameter set, we estimated the diffusion constant *D*_0_ of a DNA segment in our simulations by experimentally tracking a chromosomal locus. The origin was tagged using the same *parS*/ParB fluorescent operator/repressor system (FROS) as discussed above. For N-P_*cadBA*_-*parS_ori* and Δ*cadC-parS_ori*, a wild type *E. coli* and an *E. coli* strain lacking *cadC* each containing *parS_ori,* fluorescence and phase contrast microscopy time lapse videos were taken of the same cells every 30 s for activating and inactivating conditions, respectively.

The mean square displacement (MSD) was obtained by selecting the closest spots in subsequent image frames and calculating the ensemble-averaged MSD, the result of it being in good agreement with other tracking experiments of chromosomal loci in *E. coli* [39,40] (see S2 Fig). Previous experiments showed that DNA diffusion in *E. coli* agrees well with the Rouse model [41], reporting diffusion exponents in the range 0.4-0.6 [39,40,42]. We therefore fitted the mean square displacement to MSD(*τ*) = Γ*τ*^0.5^, obtaining Γ = 0.0111 ± 0.0001 μm^2^ s^-0.5^ for *E. coli* wild type under activating conditions and Γ = 0.0091 ± 0.0001 μm^2^ s^-0.5^ for Δ*cadC* under inactivating conditions. The DNA mobility seems to be independent of the probed conditions, since the two values do not differ significantly. Hence we used the average value 〈*τ*〉 to compute the diffusion constant *D*_0_ of a Kuhn segment according to the Rouse model.

The calculated diffusion constant of *D*_0_ ≈ 0.006 μm^2^ s^-1^ was used to scale the mean search time in Fig 4C. For the plateau value we obtained 〈*τ*_plateau_〉 ≈ 3.66 ± 0.04 min, which is surprisingly close to the experimentally measured values, considering the simplicity of our simulations. Even without consideration of non-specific binding and sliding the value only deviates from the experimental measurement by a factor ~ 1.2.

### Simulating multiple DNA-binding sites

Motivated by the experiments with the N+T-P_cadB_A strain, we performed simulations with varying number of DNA-binding sites. In Fig 4D the mean search time 〈*τ*〉 normalized by the mean search time with a single DNA-binding site 〈*τ*〉_1_ is shown as a function of the number of binding sites *N*_b_. Placing the binding sites as far apart as possible on the circular chromosome, simulations with realistic parameters show a halving in search time when increasing the number of binding sites from one to two. This is the expected result for two binding sites moving independently of each other due to the decorrelation of polymer subchains in spatial confinement. How far two binding sites have to be apart along the DNA to behave as independent targets therefore depends on the size of the simulated cell. The initial inverse-N_b_ scaling of 〈*τ*〉 flattens progressively as the binding sites come closer to one another and are more correlated in their movement. Placing the binding sites next to each other (orange stars) has an expectedly small effect for small *N*_b_, since it only increases the size of the binding site. The curve becomes steeper as the binding sites occupy a larger fraction of the polymer.

Given that the two DNA-binding sites in N+T-P_*cadBA*_ are on opposite sides of the chromosome, we expect them to behave as two independent binding sites. The experimentally measured reduction of the search time by roughly a factor of two is therefore in good agreement with our model. Also, by construction, the position of the binding site along the chromosome has no effect on the simulated search times.

### CadC diffusion strongly affects the target search time

To address the question whether DNA mobility or CadC diffusion is more crucial for the search process, numerical simulations with varying diffusion rates were run. Plotting the mean search time as a function of the CadC diffusion rate *k*_2D_ in Fig 4E we observe three different regimes. For slow protein diffusion the search time is almost independent of *k*_2D_, it follows a range where the search process is entirely dominated by CadC diffusion 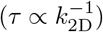 and for very fast CadC diffusion it becomes less critical again. The data point for infinitely fast CadC diffusion was obtained by making CadC cover the whole cell surface. The realistic value for the diffusion constant of CadC (*k*_2D_ = 0.20 μm^2^ s^-1^) is marked by a gray dashed line and lies in the regime where the search time strongly depends on *k*_2D_. Fig 4F shows the counterpart of this plot for DNA diffusion, marking *k*_DNA_ = 0.0060 μm^2^ s^-1^ with a gray dashed line. While the mean search time is strongly dependent on DNA diffusion for small and large *k*_DNA_, it is almost constant in the experimentally relevant intermediate regime.

To further validate this observation, we used our numerical simulations to approximate a target search exclusively due to CadC diffusion in the membrane. We used the realistic parameters but placed the DNA-binding site at the membrane and set the DNA diffusion rate to zero, such that only CadC was moving, yielding a mean search time of 〈*τ*〉 ≈ 18 s. The previously reported [43] approximate formula 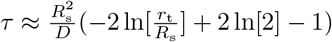 for the mean first encounter time of a particle moving with diffusion constant *D* on the surface of a sphere with radius *R*_s_ with a trap of radius *r*_t_ leads, approximating the cubic cell by a sphere of radius *R*_s_ = 0.69 μm and CadC with *r*_t_ = 0.050 μm to τ ≈ 16 s. Using the approximation does not only validate our simulations, but shows that the dependence of the search time on the size of CadC is weak, rendering a correction for overestimating the size of CadC dimers in the simulations unnecessary.

While the DNA is likely to be the less mobile part in the target search, it has to move at least close to the membrane to enable binding to CadC. The data point on the very right of Fig 4E corresponds to the time it takes the DNA-binding site to bind anywhere to the membrane. It corresponds to *τ* ≈ 14 s, similar to the time it takes CadC to locate the binding site at the membrane. Therefore, a scenario where the DNA-binding site randomly reaches the membrane and CadC searches the membrane to bind to it seems to lead to reasonable search times according to our simulations.

For comparison we also simulated a target search process where CadC is immobile and DNA diffusion has to account for the whole search. When only the DNA is moving and all other rates are set to zero, with realistic values for DNA length and cell size a mean search time of 〈*τ*〉 ≈ 560 min was calculated from the simulations, a response time that would not allow *E. coli* cells to survive the transition to acidic environments. Using the approximate formula 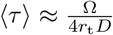 [44] for the encounter time of a particle with diffusion constant *D* to locate a target of radius *r*_t_ on the surface of a confining spherical domain with volume *V* we obtained *τ* ≈ 14 min for a monomer with diffusion constant D_o_ and *τ* ≈ 1400 min for a monomer with diffusion constant 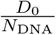. Finding a value between these two approximations is what we expected, as they do not account for the polymer dynamics.

### Quantitative comparison of simulated and experimental search time distributions

As shown above, the numerical simulations of the target search using our experimentally estimated parameter set yield mean search times that are fairly close to what we measured experimentally. To test whether the biophysical model can also quantitatively capture the experimental behavior, we attempted to find simulations with time distributions closely matching the experimentally computed CDFs. In Fig 5A the experimental CDF of N-P_*cadBA*_ together with the best fit to the sequential model with mixed initial condition is shown in orange. In order to find a simulation with agreeing time distributions, we simulated the target search process using the estimated parameter set but improved the agreement by increasing the simulated cell volume to V_cell_ = 1.3 μm^3^. As shown by the blue dots in Fig 5A this yields a very good accordance between experimental data and simulations. In Fig 5B the experimental CDF of N+T-P_*cadBA*_ and the corresponding fit to the sequential model with fixed initial condition is shown in orange. Our attempt to find a matching simulation by using the same parameters as for the simulations in panel A but increasing the DNA-binding sites to *N*_b_ = 2 leads to quite a good agreement with the experiment, as shown by the blue dots. Even though we implemented non-specific binding of CadC and sliding along the DNA, this was not necessary to obtain simulations that reproduce the experimental time distributions. Despite the simplicity of the simulations, neglecting in particular constraints of DNA movement due to its specific organization in the cell, the results agree surprisingly well with the experimental findings when using experimentally realistic parameters.

**Fig 5.**
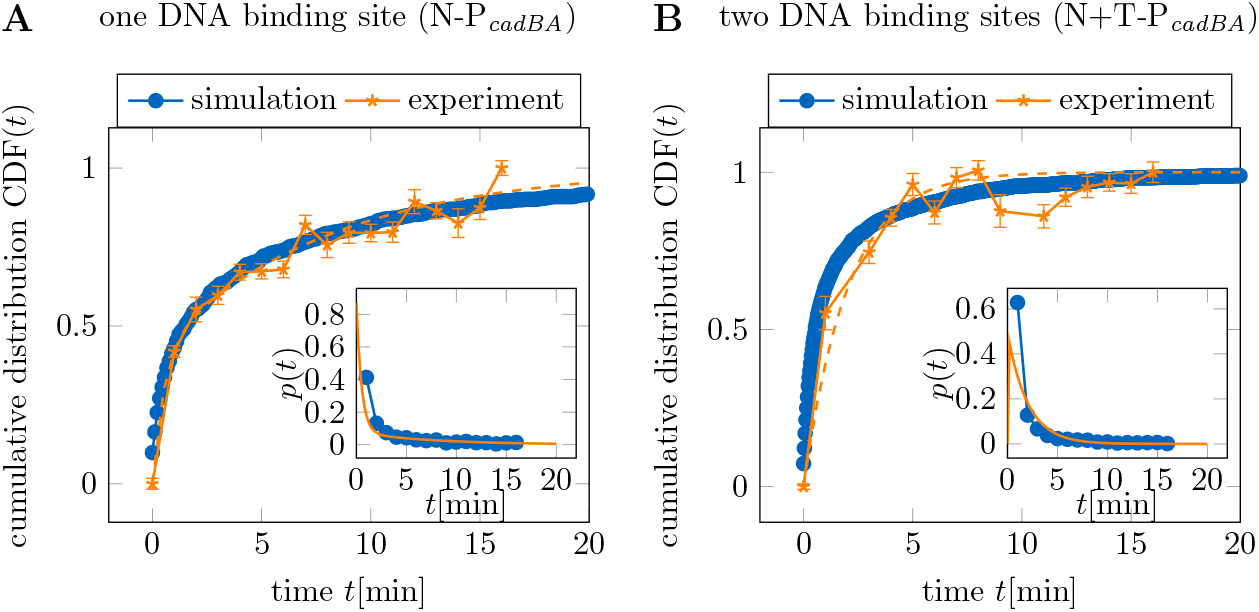
Matching simulations to the experimental results. Numerical simulations are directly compared to the experimental distributions. Experimental data are shown in orange stars, the orange dashed line corresponds to the best fit. Blue dots correspond to the simulations that agree most with the experimental data. Insets depict the corresponding first passage time distributions. (**A**) The experimental CDF of N-P_*cadBA*_ is shown together with the best fit and simulations using the realistic parameter set, summarized in Table 2. To yield better agreement with the experimental CDF the cell volume was increased to V_cell_ = 1.3 μm^3^. (**B**) The experimental CDF of N+T-P_*cadBA*_ is shown together with the best fit and simulations using the parameter set matching the experimental values, with cell volume VC_ell_ = 1.3 μm^3^ and two binding sites on the polymer. All simulations were started from a random initial configuration.

To further investigate the simulated first passage time distributions, we computed them for different parameter settings and fitted to the sequential model with fixed initial condition (Eq 9) and with mixed initial condition (Eq 10) respectively. While the shape of the first passage time distribution is independent of the system size parameters, the dynamic parameters have a big effect. For most parameter settings, including the realistic parameters, we found the best agreement of simulated CDFs with the sequential model with mixed initial condition. However, for simulations with an immobile polymer or slowly diffusing CadC, the search time distribution agrees rather with the sequential model with fixed initial condition. Since both in the experimental and simulated CDFs the delay *β* is very small compared to *α*, a fit to the fixed initial model gives virtually the same result as a single exponential CDF. When either the DNA or CadC accounts for most of the search there is only one limiting rate in the process, which can therefore be described by a single exponential distribution.

## Discussion

We combined fluorescence microscopy experiments, quantitative analysis, and kinetic Monte Carlo simulations to characterize the target search kinetics of membrane-integrated transcription factors for a specific binding site on the chromosomal DNA. We were able to measure the time between the environmental stimulus and stable DNA-binding of a membrane-integrated one-component receptor, using the pH stress-sensing receptor CadC in *E. coli* as a model system. The measured mean search time of on average 4. 5 min for a single DNA-binding site is consistent with the timescale of the earliest transcriptional response [26]. Given the severe constraint of a membrane-anchored target search, it seems surprising that the search time is only about 5-fold slower than the search time of the cytosolic Lac repressor for its operator, which takes around one minute at a similarly low protein level [12].

As the position of the DNA-binding site along the chromosome has no influence on the mean search time, the target search process appears to be quite robust. Given that the chromosome is highly organized within the cell, we believe this effect to arise mostly due to the mobility of CadC. While we do not have new evidence against proteolytic processing, this leads us to favor a diffusion and capture mechanism over transertion. Diffusion and capture mechanisms are well established for the localization of membrane-integrated proteins like SpoIVB in *Bacillus subtilis* [45,46]. As the search time is independent of the position of the DNA-binding site, transertion of CadC is at least not a requirement for fast response, in agreement with a previous evaluation of the three models [28]. Our finding that the mean search time decreases by a factor of two in a mutant with two DNA-binding sites is also consistent with the diffusion and capture mechanism. Our simulations show that this is the expected result for two independent and equally accessible binding sites, where distant parts of the polymer become uncorrelated in their motion due to the confinement in the cell.

Despite the simplicity of our biophysical model of the target search, we obtained search times that match the experimental measurements surprisingly well, even without consideration of non-specific binding and sliding. It therefore does not seem to require a fine-tuned strategy to make the target search work. Although the simulations are clearly oversimplifying the biological situation by neglecting chromosomal loci being constrained to move within macrodomains, the agreement between simulations and experimental data is striking.

While it is difficult to experimentally answer the question whether CadC or the DNA contribute more to the search process, the consistency between our simulations and experimental findings allows to draw some conclusions from an analysis of the simulations. Evaluating the functional dependence of the mean search time on the diffusion rate of the CadC dimer, the search time was found to be inversely proportional to the diffusion rate of CadC, but is almost independent of the DNA diffusion rate in the experimentally relevant regime. The search process therefore seems to be predominantly limited by the mobility of the transcription factors.

Beyond mean values the experimental data allowed the extraction of the cumulative distribution functions (CDF) for the target search time. Describing them by the distribution obtained from a reversible sequential two-step process resulted in a good fit for the wild type data assuming a mixed initial condition. The same was true for our simulations of the target search process with most parameter settings. Using a parameter set estimated for the experimental conditions led to a distribution that matched the experimental CDF of *E. coli* wild type. Attempting to find a matching simulation for the mutant with two DNA-binding sites by using the same parameters with two DNA-binding sites also yielded a good agreement. In contrast to the simulations, however, the experimental CDF of the mutant with two binding sites and the mutant with the binding site at *ter* show a better agreement with the sequential model with fixed initial condition (or a single-exponential CDF). How exactly the position of the DNA-binding site affects the initial condition of the search process is not clear at this point. In the simulations, distributions agreeing with the sequential model with fixed initial conditions are obtained when either only CadC is moving, or it is moving very slowly.

Taken together, our experiments and simulations indicate that CadC is highly mobile in the membrane, while the *cadBA* promoter on the *E. coli* chromosome is mobile enough to randomly reach the membrane, enabling CadC to locate the DNA-binding site within about five minutes, independent of its position along the chromosome. While diffusion and capture mechanisms are established for the polar localization of membrane proteins [45–47], our study indicates a broader relevance of diffusion and capture mechanisms for the largely uncharacterized interactions of membrane-integrated proteins with chromosomal DNA [23].

## Materials and methods

### Model and simulations

Since the target search of CadC for its DNA-binding site(s) has been experimentally shown to succeed within a few minutes, we were aiming at finding the simplest model that can reproduce this fast response. Hence we constrained our simulations to the main components of the target search: The DNA with one or more specific binding sites moving inside an *E. coli* cell and one or more CadC beads diffusing in the cell membrane. CadC dimerization was not simulated explicitly, instead the CadC beads on the cell surface correspond to already formed dimers. Besides diffusion of the DNA and CadC, non-specific binding of CadC to any of the DNA sites, as well as one-dimensional sliding along DNA segments that are aligned along the membrane were included.

To simulate a system as large as the whole genome in an *E. coli* cell for a time span long enough to capture the target search of CadC for its DNA-binding site is computationally challenging. We therefore chose a coarse-grained cubic lattice model in order to reduce the simulation time. In the simple cubic lattice model, the cell is composed of 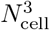 lattice points forming a cuboid. The lattice constant was chosen equal to the Kuhn length *l*_k_ = 2*l*_p_ of double stranded DNA (persistence length *l*_p_ ≈ 50 nm [48]), such that the polymer could be represented as a freely jointed chain (FJC) on the lattice. It is composed of *N*_DNA_ beads with coordinates *r* = (*r*_1_,..., *rN*_DNA_) placed on the grid points connected by *N*_DNA_ — 1 bonds, one or more of the beads being specified as the specific binding sites. Each CadC dimer occupies a single lattice point on the cell surface. In analogy to the single-site Bond-Fluctuation model [49] the polymer beads are connected by bonds of variable length, that are free to occupy also diagonal configurations. As we simulate a phantom chain, we replace the upper and lower bound of the bond length intended to ensure excluded volume and prevent chain crossings by a spring potential

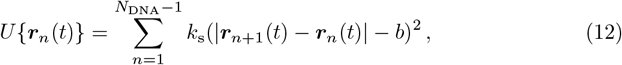

with spring constant *k*_s_ and an equilibrium bond length *b* = *l*_k_. Following the Bond-Fluctuation model, the polymer is moved by attempting the displacement of a single bead to one of the nearest lattice points, the spring potential being enforced by a Metropolis algorithm [50]. CadC beads are also moved by random displacements but constrained to the cell surface or to the DNA when bound. The kinetic Monte Carlo algorithm simulates the master equation

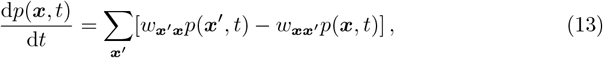

with *p*(***x***, *t*) the probability that the system is in state ***x*** at time t and transition rates *w_**x’x**_* = *w*(***x’ x***) between the states. The rates governing the dynamics are the polymer diffusion rate *k*_DNA_, which is set to one unless otherwise stated, the rate for CadC diffusion on the cell surface *k*_2D_, the non-specific binding rate *k*_on_, the unbinding rate *k*_off_ and the one-dimensional sliding rate of CadC along the polymer *k*_1D_. Unless otherwise stated the simulations were initialized with a random walk configuration of the polymer inside the cell and a randomly placed CadC bead in the membrane. To compute mean values and search time distributions ≥ 10^3^ simulations were run for every choice of parameters, each starting from a different initial configuration.

For quantitative comparison with the experiments, we defined a parameter set that matches the experimental values, as summarized in Table 2. The *E. coli* cell volume of ∝ μm^3^ is approximated by a simulation box of volume 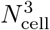 with *N*_cell_ = 10*a* and *a* = 100 nm. The *E. coli* chromosome with 4 639 221 bp, which measures 1.58 mm corresponds to a length of 15773a. Since our simulations have shown that the target search dynamics are independent of polymer length once it has reached a cell size dependent critical length (see Fig 4C), we save computation time by choosing the polymer length well above this threshold at 100a. For the CadC dimer diffusion a value of *D*_cadC_ ≈ 0.20 μm^2^ s^-1^ agrees with both previous measurements of a membrane-integrated protein with two transmembrane domains in *E. coli* [51] and single-molecule measurements of CadC [28]. From our *ori* tracking experiments we calculated the diffusion constant of a DNA bead in our simulation to be *D*_0_ ≈ 0.0060 μm^2^ s^-1^, which is the value we chose for the realistic parameter set. To convert the dimensionless search times τ’ from the simulations into seconds we computed 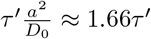.

**Table 2.** Simulation parameters matching the experiments.

| Parameter | Symbol | Value |
| --- | --- | --- |
| lattice constant | $a$ | 100 nm |
| cell volume | $V_{\text{cell}}$ | $1 \mu\text{m}^3$ |
| DNA length | $N_{\text{DNA}}$ | $10 \mu\text{m}$ |
| number of CadC dimers | $N_{\text{p}}$ | 1 |
| number of binding sites | $N_{\text{b}}$ | 1 |
| CadC diffusion | $k_{2\text{D}}$ | $0.20 \mu\text{m}^2 \text{s}^{-1}$ |
| DNA diffusion | $k_{\text{DNA}}$ | $0.0060 \mu\text{m}^2 \text{s}^{-1}$ |

Parameter set for the kinetic Monte Carlo simulations that were chosen to match the experiments.

### Experiments

#### Construction of strains and plasmids

Molecular methods were carried out according to standard protocols or according to the manufacturer’s instructions. Kits for the isolation of plasmids and the purification of PCR products were purchased from Süd-Laborbedarf (SLG; Gauting, Germany). Enzymes were purchased from New England BioLabs (Frankfurt, Germany). Bacterial strains and plasmids used in this study are summarized in table 1.

*E. coli* strains were cultivated in LB medium (10gl^-1^ NaCl, 10gl^-1^ tryptone, 5gl^-1^ yeast extract) or in Kim Epstein (KE) medium [52] adjusted to pH 5.8 or pH 7.6, using the corresponding phosphate-buffer. *E. coli* strains were always incubated aerobically in a rotary shaker at 37 °C. KE medium was always supplemented with 0.20%(w/v) glucose. Generally, lysine was added to a final concentration of 10 mmol unless otherwise stated. If necessary, media were supplemented with 100 µg ml^-1^ ampicillin or 50 μg ml^-1^ kanamycin sulfate. To allow the growth of the conjugation strain *E. coli* WM3064, we added meso-diamino-pimelic acid (DAP) to a final concentration of 200 μmol.

In order to gain strain *E. coli* MG1655-*parS_ori,* the *parS* site of *Yersinia pestis* was inserted at the origin of replication (ori) at 84.3’ in *E. coli* MG1655. Briefly, the *parS* region was inserted between *pstS* and *glmS*. Therefore, DNA fragments comprising 650 bp of *pstS* and *glmS* and the *parS’* region were amplified by PCR using MG1655 genomic DNA as template and the plasmid pFH3228, respectively. After purification, these fragments were assembled via Gibson assembly [53] into EcoRV-digested pNPTS138-R6KT plasmid, resulting in the pNTPS138-R6KT-parS_ori plasmid. The resulting plasmid was introduced into *E. coli* MG1655 by conjugative mating using *E. coli* WM3064 as a donor on LB medium containing DAP. Single-crossover integration mutants were selected on LB plates containing kanamycin but lacking DAP. Single colonies were then streaked out on LB plates containing 10%(wt/vol) sucrose but no NaCl to select for plasmid excision. Kanamycin-sensitive colonies were then checked for targeted insertion by colony PCR and sequencing of the respective PCR fragment. In order to gain strain *E. coli* MG1655-Δ*cadC parS_ori,* the *parS* site of *Y. pestis* was inserted at the origin of replication (ori) at 84.3’ in *E. coli* MG1655-Δ*cadC* as described above.

In order to gain strain *E. coli* MG1655_P_*cadBA*_-terminus, the *cadBA* promoter region was inserted at the terminus (33.7Δ) in *E. coli* MG1655. Construction of this strain was achieved via double homologous recombination using the pNTPS138-R6KT-P_*cadBA*_-terminus plasmid [28] as described above. Correct colonies were then checked for targeted insertion by colony PCR and sequencing of the respective PCR fragment.

Details of the strains and plasmids used in this study are summarized in Table 3.

**Table 3.**
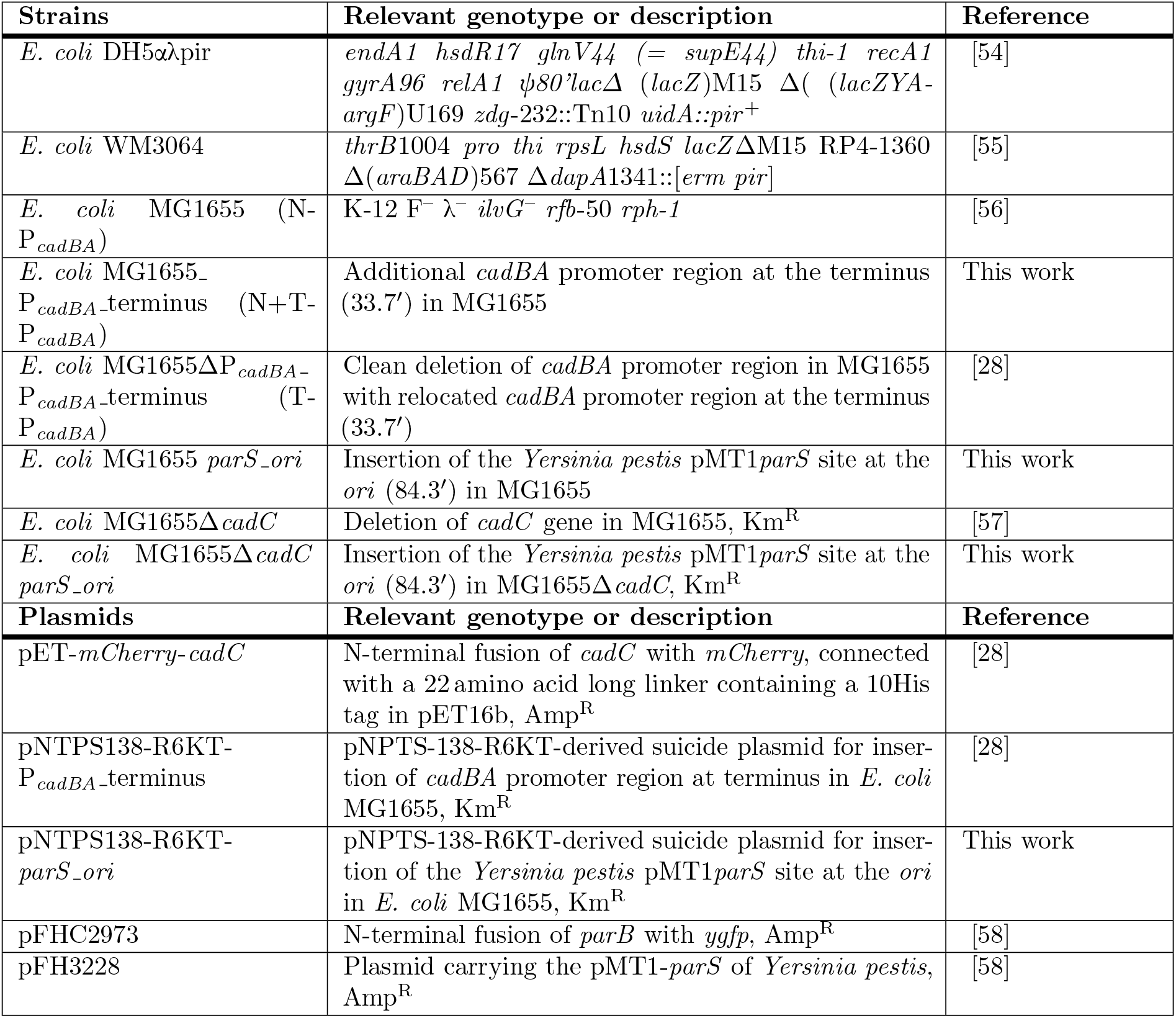
Strains and plasmids used in this study.

#### *In vivo* fluorescence microscopy

To analyze search response of mCherry-CadC to its binding site(s), overnight cultures of *E. coli* MG1655 (one CadC binding site close to ori, N-P_*cadBA*_), *E. coli* MG1655ΔP_*cddBA*_-P_*cadBA*_-terminus (one CadC binding site at terminus, T-P_*cadBA*_) and *E. coli* MG1655_P_*cadBA*_-terminus (two CadC binding sites, N+T-P_*cadBA*_), each carrying *pET-mCherry-cadC*, were prepared in KE medium pH 7.6 and aerobically cultivated at 37 °C. The overnight cultures were used to inoculate day cultures (OD_600_ of 0.1) in fresh medium at pH 7.6. At an OD_600_ of 0.5, cells were gently centrifuged and resuspended, thereby exposing them to low pH (KE medium pH 5.8 + lysine). Then the cultures were aerobically cultivated at 37°C and every 1 min after the shift to low pH, 2 μl of the culture was spotted on 1 %(w/v) agarose pads (prepared with KE medium pH 5.8 + lysine), placed onto microscope slides and covered with a coverslip. Subsequently, images were taken on a Leica DMi8 inverted microscope equipped with a Leica DFC365 FX camera (Wetzlar, Germany). An excitation wavelength of 546nm and a 605 nm emission filter with a 75 nm bandwidth was used for mCherry fluorescence with an exposure of 500 ms, gain 5, and 100 % intensity. Before shifting the cells to low pH, 2 µl of the cultures in KE medium pH 7.6 were spotted on 1 %(w/v) agarose pads (prepared with KE medium pH 7.6) and imaged as a control.

To analyze the spatiotemporal localization of a chromosomal locus, the *parS* site was inserted close to the *ori*. The localization of the *parS* site was visualized via the binding of ParB-yGFP [58]. *E. coli* MG1655 *parS~ori* cells carrying plasmid pFH3228 were cultivated in KE medium pH 7.6 as described above. At an OD_600_ of 0.5, 2 µl of the culture were shifted on 1 %(w/v) agarose pads (prepared with KE medium pH 7.6 or pH 5.8 + lysine) and placed onto microscope slides and covered with a coverslip. Subsequently, every 30 s time lapse images of the same cells were taken on a Leica DMi8 inverted microscope equipped with a Leica DFC365 FX camera (Wetzlar, Germany) of the same positions. An excitation wavelength of 485 nm and a 510 nm emission filter with a 75 nm bandwidth were used for ParB-yGFP fluorescence with an exposure of 350 ms, gain 3, and 100 % intensity.

#### Microscopy image analysis

To analyze the fluorescence microscopy images for CadC or ParB spots within the cells, we used Oufti [59], an open-source software designed for the analysis of microscopy data for cell segmentation of the phase contrast microscopy images. The resulting cell outlines were used in a custom-written software implemented in Matlab and available on request to detect fluorescent spots. Briefly, a graphical user interface (GUI) was implemented that allows testing the parameters in a test mode before running the actual detection. In detection mode a function *SpotDetection.m* is called, that iterates through all frames and all cells. For each cell, from pixels in the fluorescence microscopy images the intensity of which is above a threshold defined by the parameters and dependent on the mean and variance of the fluorescence signal within the cell the connected components are computed. The components are checked for minimum and maximum size and minimum distance to other spots before being added to the list of spots. For further computations, information on all cells and spots were saved for all frames corresponding to a certain time after receptor activation.

#### Interpretation of CadC spot data

In a custom-written Matlab script the results from the image analysis were used to compute the fraction of cells with spots *v*(*t*) as a function of time *t* after receptor activation, which upon normalization corresponds to the CDF of the search time distribution. We fit the data to the CDF of a theoretical model using the *curve_fit* function of the scipy module in Python, choosing a trust region reflective algorithm, which is an evolution of the Levenberg-Marquardt method that can handle bounds. This algorithm was developed to solve nonlinear least squares problems and combines the gradient descent method and the Gauss-Newton method. It minimizes the sum of the weighted squares of the errors between the measured data *y_i_* and the curve-fit function 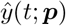

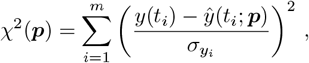

with an independent variable *t* and a vector of *n* parameters ***p*** and a set of m data points (*t_i_,y_i_*). *σ_yi_* is the measurement error for measurement *y*(*t_i_*) and 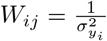 is the weighting matrix. [60]

#### Calculation of the ParB diffusion constant

The results from the image analysis were used to compute trajectories of ParB spots in a custom-written Matlab script by selecting the closest spots in subsequent image frames.

From the trajectories of ParB spots the ensemble-averaged mean square displacement (MSD) was computed as a function of time lag 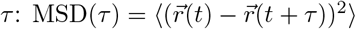, where the mean was taken over different spots and *τ* = *n*30s with *n* ∈ {1,..., *N*} and the number of time steps *N*. The Rouse model predicts the MSD in 2D [61]:

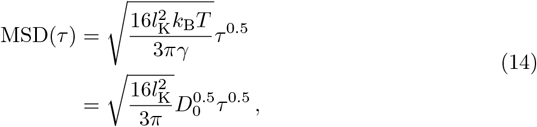

with Boltzmann’s constant *k*_B_, absolute temperature *T*, Kuhn length *l*_K_, friction constant *γ* and bead diffusion constant *D*_0_, where we have used 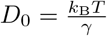 [62]. We fitted the experimentally determined MSD to MSD(*t*) = Γ*t*^0.5^ and determined the diffusion constant from 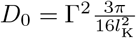.

## Supporting information

**S1 Fig. Dynamics of the target search by CadC.** Fluorescent microscopy images were taken every minute after receptor activation and analyzed for CadC spots for all three *E. coli* strains. The plot shows the fraction of cells with spots *v*(*t*) as a function of time t after the medium shift to low pH and lysine.

**S2 Fig. Mean square displacement of ParB spots.** The MSD of ParB spots was calculated by selecting the closest spots in subsequent image frames and calculating the ensemble-averaged mean square displacement as a function of time lag *τ*. The dashed lines show the fit to Γ*τ^α^*. For each time lag the mean was taken over 234 to 936 values.

**S1 Table. Fit results.** Results from fitting the experimentally computed CDF to the sequential reversible model with mixed initial condition (N-P_*cadBA*_) and fixed initial condition (T-P_*cadBA*_ and N+T-P_*cadBA*_). The fit parameters *α, β* and *c* were used to compute the mean first passage time and the variance with uncertainties obtained from error propagation using the full covariance matrix.

**S2 Table. Covariance matrix.** Covariance matrix of the parameters *α, β* and *c* from fitting the experimentally computed CDF to the sequential reversible model with mixed initial condition (N-P_*cadBA*_) and fixed initial condition (T-P_*cadBA*_ and N+T-P_*cadBA*_).

**S1 Appendix. Supplementary text.**

## Acknowledgments

We thank Flemming G. Hansen, Technical University of Denmark, for providing the plasmids pFHC2973 and pFH3228. This work was supported by the German Research Council (DFG) within the framework of the Transregio 174 “Spatiotemporal dynamics of bacterial cells” (to K.J. and U.G.).

## References

1. Hall-Stoodley L, Costerton JW, Stoodley P. Bacterial biofilms: from the natural environment to infectious diseases. Nature reviews microbiology. 2004;2(2):95.

2. Ulrich LE, Koonin EV, Zhulin IB. One-component systems dominate signal transduction in prokaryotes. Trends in microbiology. 2005;13(2):52–56.

3. Stock AM, Robinson VL, Goudreau PN. Two-component signal transduction. Annual review of biochemistry. 2000;69(1):183–215.

4. Riggs AD, Bourgeois S, Cohn M. The Lac repressor-operator interaction: III. Kinetic studies. Journal of molecular biology. 1970;53(3):401–417.

5. Adam G, Delbrück M. Reduction of dimensionality in biological diffusion processes. Structural chemistry and molecular biology. 1968;198:198–215.

6. Richter PH, Eigen M. Diffusion controlled reaction rates in spheroidal geometry: application to repressor-operator association and membrane bound enzymes. Biophysical chemistry. 1974;2(3):255–263.

7. Berg OG, Winter RB, Von Hippel PH. Diffusion-driven mechanisms of protein translocation on nucleic acids. 1. Models and theory. Biochemistry. 1981;20(24):6929–6948.

8. Slutsky M, Mirny LA. Kinetics of protein-DNA interaction: facilitated target location in sequence-dependent potential. Biophysical Journal. 2004;87(6):4021–4035.

9. Hu T, Grosberg AY, Shklovskii B. How proteins search for their specific sites on DNA: the role of DNA conformation. Biophysical journal. 2006;90(8):2731–2744.

10. Bonnet I, Biebricher A, Porte PL, Loverdo C, Bénichou O, Voituriez R, et al. Sliding and jumping of single EcoRV restriction enzymes on non-cognate DNA. Nucleic acids research. 2008;36(12):4118–4127.

11. Loverdo C, Benichou O, Voituriez R, Biebricher A, Bonnet I, Desbiolles P. Quantifying hopping and jumping in facilitated diffusion of DNA-binding proteins. Physical review letters. 2009;102(18):188101.

12. Elf J, Li GW, Xie XS. Probing transcription factor dynamics at the single-molecule level in a living cell. Science. 2007;316(5828):1191–1194.

13. Hammar P, Leroy P, Mahmutovic A, Marklund EG, Berg OG, Elf J. The Lac repressor displays facilitated diffusion in living cells. Science. 2012;336(6088):1595–1598.

14. van den Broek B, Lomholt MA, Kalisch SM, Metzler R, Wuite GJ. How DNA coiling enhances target localization by proteins. Proceedings of the National Academy of Sciences. 2008;105(41):15738–15742.

15. Schötz T, Neher RA, Gerland U. Target search on a dynamic DNA molecule. Physical Review E. 2011;84(5):051911.

16. Li GW, Berg OG, Elf J. Effects of macromolecular crowding and DNA looping on gene regulation kinetics. Nature Physics. 2009;5(4):294.

17. Liu L, Cherstvy AG, Metzler R. Facilitated diffusion of transcription factor proteins with anomalous bulk diffusion. The Journal of Physical Chemistry B. 2017;121(6):1284–1289.

18. Miller VL, Taylor RK, Mekalanos JJ. Cholera toxin transcriptional activator ToxR is a transmembrane DNA binding protein. cell. 1987;48(2):271–279.

19. Shaner NC, Lambert GG, Chammas A, Ni Y, Cranfill PJ, Baird MA, et al. A bright monomeric green fluorescent protein derived from *Branchiostoma lanceolatum*. Nature methods. 2013;10(5):407.

20. Yang Y, Isberg RR. Transcriptional regulation of the *Yersinia pseudotuberculosis* pH 6 antigen adhesin by two envelope-associated components. Molecular microbiology. 1997;24(3):499–510.

21. Buchner S, Schlundt A, Lassak J, Sattler M, Jung K. Structural and functional analysis of the signal-transducing linker in the pH-responsive one-component system CadC of *Escherichia coli*. Journal of molecular biology. 2015;427(15):2548–2561.

22. Woldringh CL. The role of co-transcriptional translation and protein translocation (transertion) in bacterial chromosome segregation. Molecular microbiology. 2002;45(1):17–29.

23. Roggiani M, Goulian M. Chromosome-membrane interactions in bacteria. Annual review of genetics. 2015;49:115–129.

24. Rodriguez MAP, Guo X. Biomacromolecular localization in bacterial cells by the diffusion and capture mechanism. Annals of Microbiology. 2013;63(3):825–832.

25. Jung K, Fabiani F, Hoyer E, Lassak J. Bacterial transmembrane signalling systems and their engineering for biosensing. Open biology. 2018;8(4):180023.

26. Fritz G, Koller C, Burdack K, Tetsch L, Haneburger I, Jung K, et al. Induction kinetics of a conditional pH stress response system in *Escherichia coli*. Journal of molecular biology. 2009;393(2):272–286.

27. Haneburger I, Fritz G, Jurkschat N, Tetsch L, Eichinger A, Skerra A, et al. Deactivation of the *E. coli* pH stress sensor CadC by cadaverine. Journal of molecular biology. 2012;424(1-2):15–27.

28. Brameyer S, Rösch TC, El Andari J, Hoyer E, Schwarz J, Graumann PL, et al. DNA-binding directs the localization of a membrane-integrated receptor of the ToxR family. Communications biology. 2019;2(1):4.

29. Görke B, Reinhardt J, Rak B. Activity of Lac repressor anchored to the *Escherichia coli* inner membrane. Nucleic acids research. 2005;33(8):2504–2511.

30. Redner S. A guide to first-passage processes. Cambridge University Press; 2001.

31. Schlundt A, Buchner S, Janowski R, Heydenreich T, Heermann R, Lassak J, et al. Structure-function analysis of the DNA-binding domain of a transmembrane transcriptional activator. Sci Rep. 2017;7(1):1051.

32. Haneburger I, Eichinger A, Skerra A, Jung K. New insights into the signaling mechanism of the pH-responsive, membrane-integrated transcriptional activator CadC of *Escherichia coli*. Journal of Biological Chemistry. 2011;286(12):10681–10689.

33. Tetsch L, Koller C, Haneburger I, Jung K. The membrane-integrated transcriptional activator CadC of *Escherichia coli* senses lysine indirectly via the interaction with the lysine permease LysP. Molecular microbiology. 2008;67(3):570–583.

34. Rauschmeier M, Schuöppel V, Tetsch L, Jung K. New insights into the interplay between the lysine transporter LysP and the pH sensor CadC in *Escherichia coli*. Journal of molecular biology. 2014;426(1):215–229.

35. Lindner E, White SH. Topology, dimerization, and stability of the single-span membrane protein CadC. Journal of molecular biology. 2014;426(16):2942–2957.

36. Ude S, Lassak J, Starosta AL, Kraxenberger T, Wilson DN, Jung K. Translation elongation factor EF-P alleviates ribosome stalling at polyproline stretches. Science. 2013;339(6115):82–85.

37. Zhou J, Rudd KE. EcoGene 3.0. Nucleic acids research. 2012;41(D1):D613–D624.

38. Youngren B, Nielsen HJ, Jun S, Austin S. The multifork *Escherichia coli* chromosome is a self-duplicating and self-segregating thermodynamic ring polymer. Genes & development. 2014;28(1):71–84.

39. Weber SC, Spakowitz AJ, Theriot JA. Bacterial chromosomal loci move subdiffusively through a viscoelastic cytoplasm. Physical review letters. 2010;104(23):238102.

40. Espeli O, Mercier R, Boccard F. DNA dynamics vary according to macrodomain topography in the *E. coli* chromosome. Molecular microbiology. 2008;68(6):1418–1427.

41. Rouse Jr PE. A theory of the linear viscoelastic properties of dilute solutions of coiling polymers. The Journal of Chemical Physics. 1953;21(7):1272–1280.

42. Javer A, Long Z, Nugent E, Grisi M, Siriwatwetchakul K, Dorfman KD, et al. Short-time movement of *E. coli* chromosomal loci depends on coordinate and subcellular localization. Nature communications. 2013;4:3003.

43. Singer A, Schuss Z, Holcman D, Eisenberg R. Narrow escape, part I. Journal of Statistical Physics. 2006;122(3):437–463.

44. Amitai A, Amoruso C, Ziskind A, Holcman D. Encounter dynamics of a small target by a polymer diffusing in a confined domain. The Journal of chemical physics. 2012;137(24):244906.

45. Thanbichler M, Shapiro L. Getting organized—how bacterial cells move proteins and DNA. Nature Reviews Microbiology. 2008;6(1):28–40.

46. Rudner DZ, Pan Q, Losick RM. Evidence that subcellular localization of a bacterial membrane protein is achieved by diffusion and capture. Proceedings of the National Academy of Sciences. 2002;99(13):8701–8706.

47. Laloux G, Jacobs-Wagner C. How do bacteria localize proteins to the cell pole? Journal of cell science. 2014;127(1):11–19.

48. Geggier S, Kotlyar A, Vologodskii A. Temperature dependence of DNA persistence length. Nucleic acids research. 2011;39(4):1419–1426.

49. Carmesin I, Kremer K. The bond fluctuation method: a new effective algorithm for the dynamics of polymers in all spatial dimensions. Macromolecules. 1988;21(9):2819–2823.

50. Metropolis N, Rosenbluth AW, Rosenbluth MN, Teller AH, Teller E. Equation of state calculations by fast computing machines. The journal of chemical physics. 1953;21(6):1087–1092.

51. Oswald F, Varadarajan A, Lill H, Peterman EJ, Bollen YJ. MreB-dependent organization of the *E. coli* cytoplasmic membrane controls membrane protein diffusion. Biophysical journal. 2016;110(5):1139–1149.

52. Epstein W, Kim BS. Potassium transport loci in *Escherichia coli* K-12. Journal of Bacteriology. 1971;108(2):639–644.

53. Gibson DG, Young L, Chuang RY, Venter JC, Hutchison III CA, Smith HO. Enzymatic assembly of DNA molecules up to several hundred kilobases. Nature methods. 2009;6(5):343.

54. Miller VL, Mekalanos JJ. A novel suicide vector and its use in construction of insertion mutations: osmoregulation of outer membrane proteins and virulence determinants in *Vibrio cholerae* requires *toxR*. Journal of bacteriology. 1988;170(6):2575–2583.

55. Dehio C, Meyer M. Maintenance of broad-host-range incompatibility group P and group Q plasmids and transposition of Tn5 in *Bartonella henselae* following conjugal plasmid transfer from *Escherichia coli*. Journal of bacteriology. 1997;179(2):538–540.

56. Blattner FR, Plunkett G, Bloch CA, Perna NT, Burland V, Riley M, et al. The complete genome sequence of *Escherichia coli* K-12. science. 1997;277(5331):1453–1462.

57. Kuöper C, Jung K. CadC-mediated activation of the *cadBA* promoter in *Escherichia coli*. Journal of molecular microbiology and biotechnology. 2005;10(1):26–39.

58. Nielsen HJ, Ottesen JR, Youngren B, Austin SJ, Hansen FG. The *Escherichia coli* chromosome is organized with the left and right chromosome arms in separate cell halves. Molecular microbiology. 2006;62(2):331–338.

59. Paintdakhi A, Parry B, Campos M, Irnov I, Elf J, Surovtsev I, et al. Oufti: an integrated software package for high-accuracy, high-throughput quantitative microscopy analysis. Molecular microbiology. 2016;99(4):767–777.

60. Seiler MC, Seiler FA. Numerical recipes in C: the art of scientific computing. Risk Analysis. 1989;9(3):415–416.

61. Socol M, Wang R, Jost D, Carrivain P, Vaillant C, Le Cam E, et al. Rouse model with transient intramolecular contacts on a timescale of seconds recapitulates folding and fluctuation of yeast chromosomes. Nucleic acids research. 2019;.

62. Doi M, Edwards SF. The theory of polymer dynamics. vol. 73. oxford university press; 1988.

