## Supplementary Figures, Table, and Text for "Dynamics of chromosomal target search by a membrane-integrated one-component receptor"

### List of contents


- **Figure S1:** Dynamics of the target search by CadC
- **Figure S2:** Mean square displacement of ParB foci
- **Table S1:** Fit results
- **Table S2:** Covariance matrix
- **Supplementary text:**
  - Sequential reversible process
  - Mixed initial condition
  - Moments

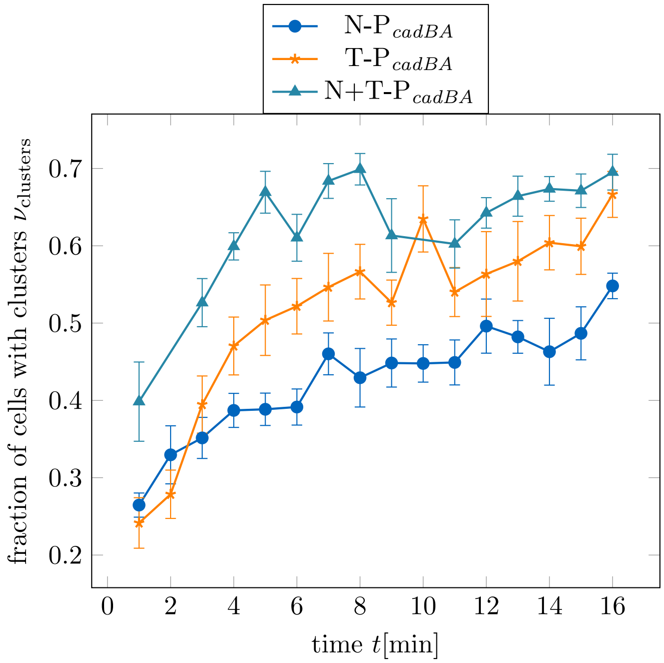


Figure 1: **Dynamics of the target search by CadC.** Fluorescent microscopy images were taken every minute after receptor activation and analyzed for CadC clusters for all three *E. coli* strains. The plot shows the fraction of cells with clusters $\nu(t)$ as a function of time $t$ after the medium shift to low pH and lysine.

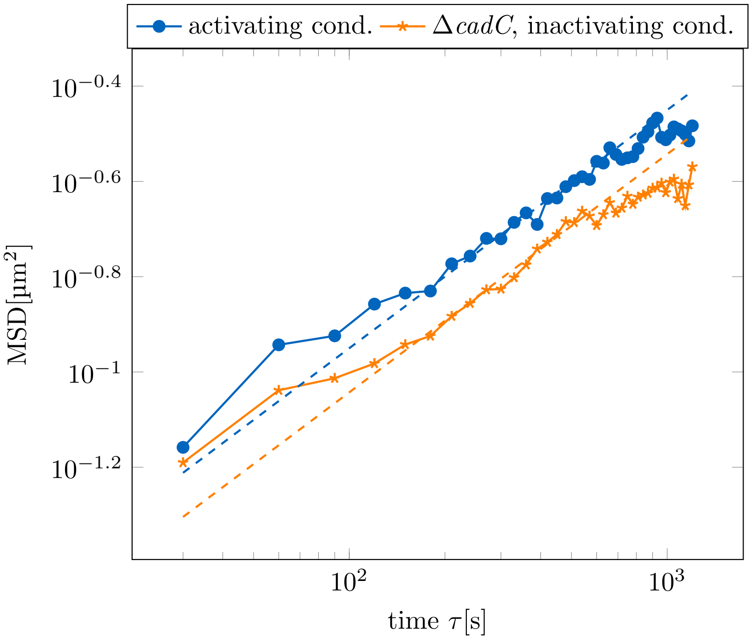


Figure 2: **Mean square displacement of ParB foci.** The MSD of ParB foci was calculated by selecting the closest foci in subsequent image frames and calculating the ensemble-averaged mean square displacement as a function of time lag $\tau$. The dashed lines show the fit to $\mathrm{MSD}(\tau) = \Gamma\tau^{\frac{1}{2}}$. For each time lag the mean was taken over 234 to 936 values.

Table 1: **Fit results.**

| Strain | $\alpha$[min] | $\beta$[min] | $c$[min$^{-1}$] | $\langle\tau\rangle$[min] | $\sigma^2$[min$^2$] |
|---|---|---|---|---|---|
| N-P$_{cadBA}$ | 7.87 ± 0.60 | 0.52 ± 0.13 | 0.87 ± 0.15 | 4.84 ± 0.19 | 49.6 ± 8.8 |
| T-P$_{cadBA}$ | 4.20 ± 0.26 | $6 \times 10^{-15}$±0.18 | | 4.20 ± 0.15 | 17.6 ± 2.0 |
| N+T-P$_{cadBA}$ | 2.02 ± 0.16 | $1.1 \times 10^{-14}$±0.22 | | 2.02 ± 0.12 | 4.09 ± 0.66 |

Results from fitting the experimentally computed CDF to the sequential reversible model with mixed initial condition (N-P$_{cadBA}$) and fixed initial condition (T-P$_{cadBA}$ and N+T-P$_{cadBA}$). The fit parameters $\alpha$, $\beta$ and $c$ were used to compute the mean first passage time and the variance with uncertainties obtained from error propagation using the full covariance matrix.

Table 2: **Covariance matrix.**

| Strain | $\sigma_{\alpha}^2$[min$^2$] | $\sigma_{\beta}^2$[min$^2$] | $\sigma_{c}^2$[min$^2$] | $\sigma_{\alpha\beta}$[min$^2$] | $\sigma_{\alpha c}$[min$^2$] | $\sigma_{\beta c}$[min$^2$] |
|---|---|---|---|---|---|---|
| N-P$_{cadBA}$ | 0.36 | 0.016 | 0.023 | 0.05 | −0.019 | −0.05 |
| T-P$_{cadBA}$ | 0.059 | 0.032 | | −0.033 | | |
| N+T-P$_{cadBA}$ | 0.026 | 0.050 | | −0.030 | | |

Covariance matrix of the parameters $\alpha$, $\beta$ and $c$ from fitting the experimentally computed CDF to the sequential reversible model with mixed initial condition (N-P$_{cadBA}$) and fixed initial condition (T-P$_{cadBA}$ and N+T-P$_{cadBA}$).

### SUPPLEMENTARY TEXT

#### Sequential reversible process

We consider a sequential reversible two-step process

$$S_1 \underset{k_2^-}{\overset{k_1^+}{\rightleftharpoons}} S_2 \overset{k_2^+}{\rightarrow} S_3\,.$$

First we consider the general case of $N$ states $n = \{1, \ldots, N\}$. The Master equation for the transition probability $p(n, t|n', t')$ from state $n'$ at time $t'$ to state $n$ at time $t$ reads

$$\frac{\partial}{\partial t} p(n, t|n', t') = k_{n-1}^+ p(n-1, t|n', t') + k_{n+1}^- p(n+1, t|n', t') - (k_n^+ + k_n^-) p(n, t|n', t')\,.$$

Since we are interested in the time $\tau$ to reach the final state $S_N$, we set $k_N^- = 0$, such that $S_N$ becomes an absorbing state. Define $G(m, t)$, the probability that at time $t$ state $S_N$ is not yet reached when starting from state $m$ at time $t = 0$:

$$G(m, t) := \mathbb{P}\{\tau \geq t|n(0) = m\} = \sum_{n=1}^{N} p(n, t|m, 0)\,.$$

The distribution of first passage times is obtained from $G(m, t)$, since

$$1 - G(m, t) = \mathbb{P}\{\tau < t|n(0) = m\}\,.$$

$1 - G(1, t)$ is the cumulative distribution function (CDF) of the first passage time, therefore

$$f(\tau) = -\frac{\partial}{\partial t} G(1, t)|_{t=\tau} \tag{1}$$

is the first passage time distribution. We exploit that the transition rates do not depend on time to write

$$G(m, t) = \sum_{n=1}^{N} p(n, 0|m, -t)\,.$$

Therefore $G(m, t)$ follows from the evolution of $p$ in the coordinate of the initial state, described by the backward Master Equation

$$\frac{\partial}{\partial t'} p(n, t|n', t') = k_{n'}^+ (p(n, t|n', t') - p(n, t|n'+1, t')) + k_{n'}^- (p(n, t|n', t') - p(n, t|n'-1, t'))\,,$$

which leads to

$$-\frac{\partial}{\partial t} G(m, t) = k_m^+ (G(m, t) - G(m+1, t)) + k_m^- (G(m, t) - G(m-1, t))$$

with initial condition $G(m, 0) = 1$ for $m < N$ and $G(N, 0) = 0$.

We now go back to the two-step process $N = 3$. Since $G(3, t) = 0$ we have two equations:

$$\begin{aligned}
-\frac{\partial}{\partial t} G(1, t) &= k_1^+ (G(1, t) - G(2, t)) + k_1^- G(1, t) \\
&= k_1^+ (G(1, t) - G(2, t)) \\
-\frac{\partial}{\partial t} G(2, t) &= k_2^+ (G(1, t) - G(3, t)) + k_2^- (G(2, t) - G(1, t)) \\
&= k_2^+ G(1, t) + k_2^- (G(2, t) - G(1, t))
\end{aligned}$$

To solve this initial value problem, we use the Laplace transform

$$\begin{aligned}
F(s) &= \int_0^\infty f(t) e^{-st} \mathrm{d}t \\
f(t) &= \frac{1}{2\pi i} \lim_{T\to\infty} \int_{\gamma - iT}^{\gamma + iT} e^{st} F(s) \mathrm{d}s\,.
\end{aligned}$$

Using $\mathcal{L}(f') = sF(s) - f(0)$ we obtain

$$\begin{aligned}
-sg(1,s) + G(1,0) &= k_1^+(g(1,s) - g(2,s)) \\
-sg(1,s) + 1 &= k_1^+(g(1,s) - g(2,s)) \\
-sg(2,s) + G(2,0) &= k_2^+ g(2,s) + k_2^-(g(2,s) - g(1,s)) \\
-sg(2,s) + 1 &= k_2^+ g(2,s) + k_2^-(g(2,s) - g(1,s))\,.
\end{aligned}$$

We solve for $g(1,s)$ and $g(2,s)$ and obtain

$$g(1,s) = \frac{k_2^- + k_1^+ + k_2^+ + s}{k_2^- s + (k_1^+ + s)(k_2^+ + s)}$$

$$g(2,s) = \frac{k_2^- + k_1^+ + s}{k_2^- s + (k_1^+ + s)(k_2^+ + s)}$$

and after back transformation

$$G(1,t) = \frac{e^{-(a+b)t}(-2b + e^{2at}2b + 2a + e^{2at}2a)}{4a} \tag{2}$$

and

$$G(2,t) = \frac{e^{-(a+b)t}(-2(b - k_2^+) + e^{2at}2(b - k_2^+) + 2a + e^{2at}2a)}{4a} \tag{3}$$

with $a = \frac{1}{2}\sqrt{(k_2^- + k_1^+ + k_2^+)^2 - 4k_1^+ k_2^+}$ and $b = \frac{1}{2}(k_2^- + k_1^+ + k_2^+)$. Using eq. (1) we obtain

$$f(\tau) = \frac{k_1^+ k_2^+}{2a}\left(e^{(a-b)\tau} - e^{-(a+b)\tau}\right) \tag{4}$$

$$= \frac{b^2 - a^2}{2a}\left(e^{(a-b)\tau} - e^{-(a+b)\tau}\right) \tag{5}$$

and for the CDF

$$\mathrm{CDF}(\tau) = 1 - G(1,t) = 1 - \frac{(a-b)e^{-(a+b)\tau} + (a+b)e^{(a-b)\tau}}{2a}\,.$$

Defining $\alpha = \frac{1}{a+b}$ and $\beta = \frac{1}{b-a}$, the CDF reads

$$\mathrm{CDF}(\tau) = 1 - \frac{\alpha e^{-\frac{\tau}{\alpha}} - \beta e^{-\frac{\tau}{\beta}}}{\alpha - \beta} \tag{6}$$

and the probability density function

$$p(\tau) = \frac{e^{-\frac{\tau}{\alpha}} - e^{-\frac{\tau}{\beta}}}{\alpha - \beta}\,. \tag{7}$$

When we assume that $k_2^- = 0$, it reduces to

$$\mathrm{CDF}(\tau) = 1 + \frac{k_2^+ e^{-k_1^+\tau} - k_1^+ e^{-k_2^+\tau}}{k_1^+ - k_2^+}\,. \tag{8}$$

When we assume that $k_2^+$ is rate limiting, it reduces to

$$\mathrm{CDF}(\tau) = 1 - e^{-k_2^+\tau}\,. \tag{9}$$

### Mixed initial condition

The above first passage time distribution is based on a system that starts initially in state $S_1$. When we allow a mixed initial state, with a fraction $x$ in state $S_2$ and a fraction $1-x$ in state $S_1$, we obtain

$$\begin{aligned}\mathrm{CDF}(\tau) &= \mathbb{P}\{\tau < t | n(0) = x s_2 + (1-x) s_1\} \\ &= x\mathbb{P}\{\tau < t | n(0) = s_2\} + (1-x)\mathbb{P}\{\tau < t | n(0) = s_1\} \\ &= 1 - xG(2,t) - (1-x)G(1,t)\end{aligned}$$

and

$$f(\tau) = -\frac{\partial}{\partial t}(xG(2,t) + (1-x)G(1,t))|_{t=\tau}\,.$$

With $G(1,t)$ and $G(2,t)$ given in eqs. (2) and (3) this yields

$$\mathrm{CDF}(\tau) = 1 - \frac{\alpha e^{-\frac{\tau}{\alpha}} - \beta e^{-\frac{\tau}{\beta}} - \alpha\beta c\left(e^{-\frac{\tau}{\alpha}} - e^{-\frac{\tau}{\beta}}\right)}{\alpha - \beta} \tag{10}$$

and

$$f(\tau) = \frac{e^{-\frac{\tau}{\alpha}} - e^{-\frac{\tau}{\beta}} + c\left(\alpha e^{-\frac{\tau}{\beta}} - \beta e^{-\frac{\tau}{\alpha}}\right)}{\alpha - \beta} \tag{11}$$

with $c = xk_2^+$ and

$$\begin{aligned}\alpha &= 2\left(k_1^+ + k_2^+ + k_2^- - \sqrt{(k_1^+ + k_2^+ + k_2^-)^2 - 4k_1^+ k_2^+}\right)^{-1}\,, \\ \beta &= 2\left(k_2^- + k_1^+ + k_2^+ + \sqrt{(k_2^- + k_1^+ + k_2^+)^2 - 4k_1^+ k_2^+}\right)^{-1}\,.\end{aligned}$$

### Moments

For the sequential reversible process with fixed initial condition we use the moments of the probability density function eq. (7) to compute the mean first passage time and the variance as functions of $\alpha$ and $\beta$. Since $\alpha$ and $\beta$ are not independent, we propagate the uncertainty with the whole covariance matrix $\sigma_{ij} = \mathrm{cov}[X_i, X_j] = \mathbb{E}[(X_i - \mathbb{E}[X_i])(X_j - \mathbb{E}[X_j]))]$, such that the uncertainty of a function $y(\alpha,\beta)$ reads

$$\begin{aligned}\Delta y &= \sqrt{\sum_{i=1}^{m}\left(\frac{\partial y}{\partial \sigma_{ii}}\right)^2 + 2\sum_{i=1}^{m-1}\sum_{j=i+1}^{m}\frac{\partial y}{\partial \sigma_{ii}}\frac{\partial y}{\partial x_j}\sigma_{ij}} \\ &= \sqrt{\left(\frac{\partial y}{\partial \alpha}\sigma_\alpha\right)^2 + \left(\frac{\partial u}{\partial \beta}\sigma_\beta\right)^2 + 2\frac{\partial y}{\partial \alpha}\frac{\partial y}{\partial \beta}\sigma_{\alpha\beta}}\,.\end{aligned}$$

We compute the mean (first moment $\mu = \mathbb{E}[X]$)

$$\begin{aligned}\mu &= \alpha + \beta \\ \Delta\mu &= \sqrt{\sigma_\alpha^2 + \sigma_\beta^2 + \sigma_{\alpha\beta}}\end{aligned}$$

and the variance (second central moment $\sigma^2 = \mathbb{E}[(X-\mu)^2]$)

$$\begin{aligned}\sigma^2 &= \alpha^2 + \beta^2 \\ \Delta\sigma^2 &= 2\sqrt{\alpha^2\sigma_\alpha^2 + \beta^2\sigma_\beta^2 + \alpha\beta\sigma_{\alpha\beta}}\,.\end{aligned}$$

For the sequential reversible model with mixed initial condition, we proceed as above using the probability density function in eq. (10). The uncertainty of a function $y(\alpha, \beta, c)$ now reads

$$\Delta y = \sqrt{\left(\frac{\partial y}{\partial \alpha}\sigma_\alpha\right)^2 + \left(\frac{\partial u}{\partial \beta}\sigma_\beta\right)^2 + \left(\frac{\partial y}{\partial c}\sigma_c\right)^2 + 2\frac{\partial y}{\partial \alpha}\frac{\partial y}{\partial \beta}\sigma_{\alpha\beta} + 2\frac{\partial y}{\partial \alpha}\frac{\partial y}{\partial c}\sigma_{\alpha c} + 2\frac{\partial y}{\partial \beta}\frac{\partial y}{\partial c}\sigma_{\beta c}}\,.$$

We again compute the mean

$$\begin{aligned}
\mu &= \alpha + \beta - \alpha\beta c \\
\Delta\mu &= \sqrt{\sigma_\alpha^2(1-\beta c)^2 + \sigma_\beta^2(1-\alpha c)^2 + \sigma_c^2\alpha^2\beta^2 + 2\sigma_{\alpha\beta}(1-\alpha c)(1-\beta c) - 2\alpha\beta(\sigma_{\alpha c}(1-\beta c) + \sigma_{\beta c}(1-\alpha c)})
\end{aligned}$$

and the variance

$$\begin{aligned}
\sigma^2 =& \alpha^2 + \beta^2 - \alpha^2\beta^2c^2 \\
\Delta\sigma^2 =& \Big\{4\sigma_\alpha^2\alpha^2(\beta^2c^2-1)^2 + 4\sigma_\beta^2\beta^2(\alpha^2c^2-1)^2 + \sigma_c^2(\beta^2 + \alpha^2c(1-\beta^2c^2))^2 + 8\sigma_{\alpha\beta}\alpha\beta(\alpha^2c^2-1)(\beta^2c^2-1) \\
&+ 8\sigma_{\alpha c}\alpha^3\beta^2c(\beta^2c^2-1) + 8\sigma_{\beta c}\alpha^2\beta^3c(\alpha^2c^2-1)\Big\}^{\frac{1}{2}}\,.
\end{aligned}$$
